# Honesty needs no cost: beneficial signals can be honest and evolutionarily stable

**DOI:** 10.1101/256248

**Authors:** Szabolcs Számadó

## Abstract

How and why animals communicate honestly is a key issue in biology. The role of signal cost is strongly entrenched in the maintenance in honest signalling. The handicap principle claims that honest signals have to be costly at the equilibrium and this cost is a theoretical necessity. The handicap principle further claims that signalling is fundamentally different from any other adaptation because honest signalling would collapse in the absence of cost. Here I investigate this claim in simple action-response game where signals do not have any cost, instead they have benefits. I show that such beneficial signals can be honest and evolutionarily stable. These signals can be beneficial to both high and low-quality signallers independently of the receiver’s response, yet they can maintain honest signalling just as much as costly signals. Signal cost-at or out of equilibrium-is not a necessary condition of honesty. Benefit functions can maintain honest signalling as long as the marginal cost-loss of benefit-is high enough for potential cheaters.

## 1. Introduction

The role of signal cost in the maintenance of honest signalling seems to be unassailable. While there is still an ongoing debate about exact nature of this role, all participants agree that some kind of cost is necessary to maintain the honesty of communication under conflict of interest [1-3]. Opinions and predictions diverge about who and when shall pay this cost. The handicap principle [1, 4] predicts the most visible and influential role for signal cost: signals have to be “wastefully” costly in order to be honest. This cost is a “test” and this test is absolutely necessary condition for honest signalling. Zahavi further argues that the selection for honest signalling is thus fundamentally different from other selection processes; the former he calls as “signal selection” vs. the “utilitarian” selection of the later [4, 5]. He argues while cost is an unavoidable “evil” for other adaptations, it is a necessity for signals [4, 5].

On the other hand, recent models of “costly signalling” paint a slightly different picture: The equilibrium cost for honest signallers can be zero or even negative, only potential cheaters have to pay a cost [6-9]. It also turned out that partially honest, so called “pooling” equilibria can be cost free [10, 11]. However, signal cost still seems to be an essential ingredient of honest signalling even in these models: (i) signals have a cost function and (ii) potential cheaters pay a cost for deviating from the equilibrium.

All in all, while these models challenge Zahavi’s main prediction about the role of equilibrium cost, they do not challenge the role of signal cost. Here I show that signal cost is not an essential ingredient of honest signalling: signals with benefit functions can be honest and evolutionarily stable even under conflict of interest. I call these signals as “beneficial signals” as opposed to “costly signals”. I show the existence of a fully honest (separating) equilibrium without any signal cost function at all. At this equilibrium both low and high-quality signallers benefit from the signals, that is, no one pays any cost at our out of equilibrium, yet the signalling system is honest and it is evolutionarily stable.

## 2. The Model

The model is a simple action-response game widely investigated in the biological literature [6, 7, 10-15]. It is a two-player game with a signaller and a receiver, where the receiver controls an indivisible resource. There are two types of signallers: low and high quality. Both type benefits from obtaining the resource. The receiver only benefits from transferring the resource to a high-quality individual. Signallers have an option to give a signal; in the standard literature this signal is costly. This signal may or may not be not be honest.

The receivers’ fitness (*F*_*r*_) depends both on the signaller’s quality (*a*), which can be high (*H*) or low (*L*) and on the receiver’s response (*z*), which can be up (*U*): to give the resource, or down (*D*): not to give the resource. The signaller’s fitness (*F*_*s*_) is the sum of the value of the resource (*V*), minus the cost of signalling (*C*). The resource may be more valuable to low or to high quality signallers, accordingly the value of the resource (*V*) both depends on the quality of the signaller (*a*) and on the receiver’s response (*z*). Last but not least, the cost of signalling (*C*) depends on the quality of the signaller (*a*) and on the signaller’s behaviour (*b*), which can be to signal (*S*) or not to signal (*N*). Accordingly, *F*_*r*_ and *F*_*s*_ can be written up respectively as follows:

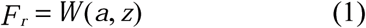

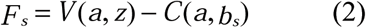

The fitness of each player can be influenced (*r*) by the survival of the other player. For example, they can be related, or they might belong to the same group (see Maynard Smith, 1991). With the help of *r* it is possible to describe different situations, for instance, where this interdependence is high (*r*>>0, e.g. parent-offspring communication) or situations without relatedness and additional interactions (i.e. *r*=0). Based on these assumptions the inclusive fitness of the signaller (*E*_*s*_) and the receiver (*E*_*r*_) can be written as follows:

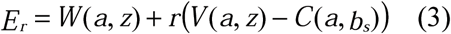

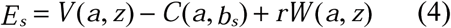

Let *V*_*h*_ and *V*_*l*_ denote the difference in fitness for high-, and low-quality signaller respectively between obtaining the resource or not (Hurd, 1995; Számadó, 1999):

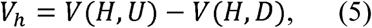

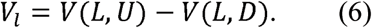

We can define *W*_*h*_, *W*_*l*_ and *C*_*h*_, *C*_*l*_ in a similar way:

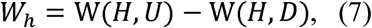

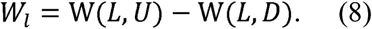

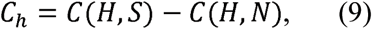

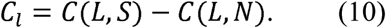

This notation will be used in the rest of the article (see Table 1. for a summary). Figure 1 depicts the signalling game, Table 2 gives the fitness values corresponding to each node.

**Table 1.** Parameters and the notation of the model.

|  |  |
| --- | --- |
| $F_r$ | receivers' fitness |
| $F_s$ | signaller's fitness |
| $W$ | value of the receiver's response for the receiver |
| $V$ | value of the receiver's response for the signaller |
| $C$ | cost of signalling |
| $a$ | signaller's quality |
| $b$ | signaller's behaviour |
| $z$ | receiver's response |
| $r$ | degree of relatedness |
| $H$ | high quality signaller |
| $L$ | low quality signaller |
| $U$ | up, to give the resource |
| $D$ | down, not to give the resource |
| $S$ | signal |
| $N$ | not to signal |
| $V_h=V(H,U)-V(H,D)$ | difference in the value of the receiver's responses for high quality signallers |
| $V_l=V(L,U)-V(L,D)$ | difference in the value of the receiver's responses for low quality signallers |
| $W_h=W(H,U)-W(H,D)$ | difference in the value of the receiver's responses for receivers in case of high quality signallers |
| $W_l=W(L,U)-W(L,D)$ | difference in the value of the receiver's responses for receivers in case of low quality signallers |
| $C_h=C(H,S)-C(H,N)$ | difference in the cost of signals for high quality signallers |
| $C_l=C(L,S)-C(L,N)$ | difference in the cost of signals for low quality signallers |

**Figure 1.**
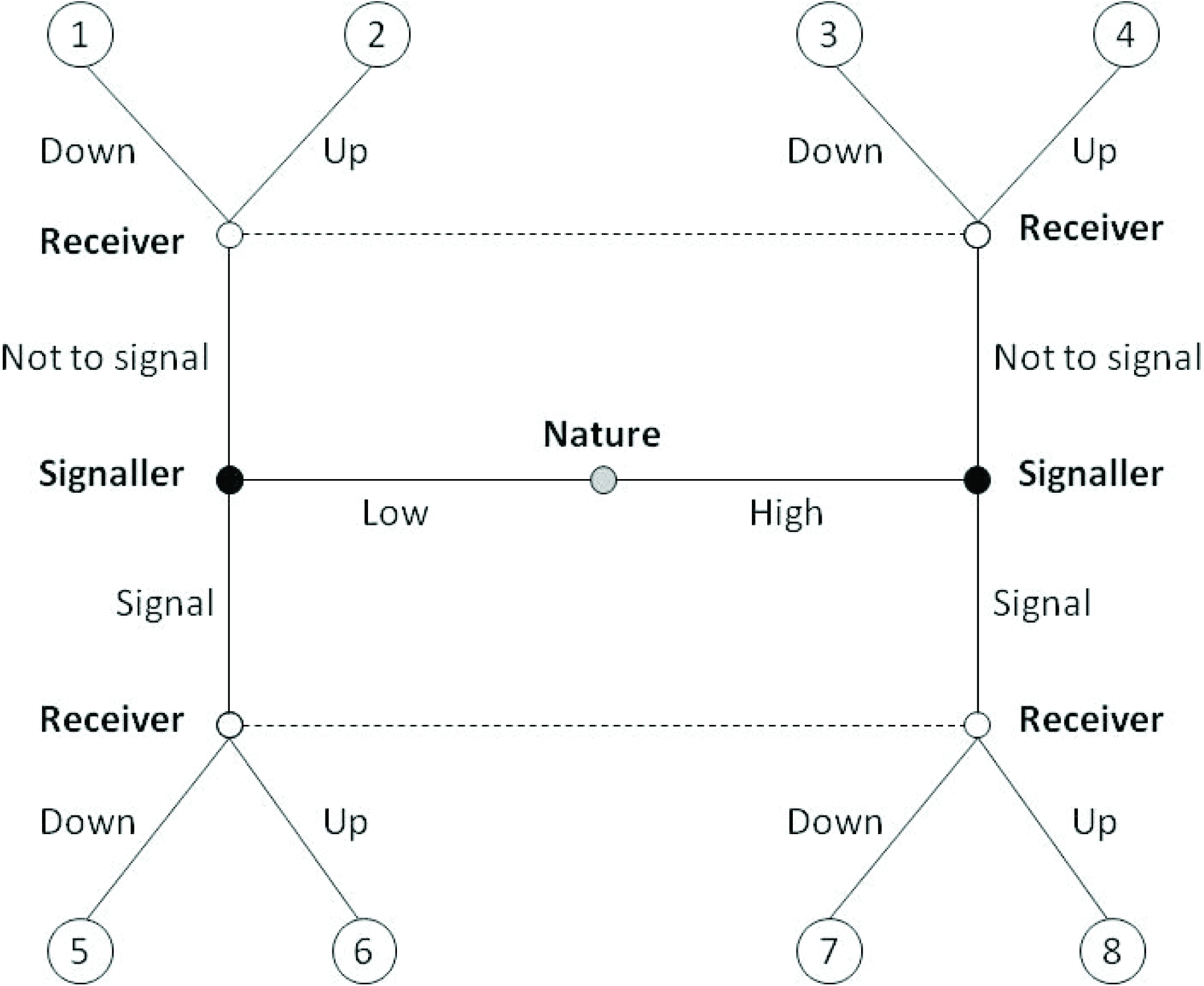
The action-response game.

**Table 2.** The fitness values corresponding to the end nodes in Figure 1, where *Es* and *Er* denote the inclusive fitness of the signaller and the receiver respectively. The fitness of both players is a combination of the benefit they receive as a result of the receiver’s decision and the costs/benefits resulting from the signaller’s decision.

| End node (Fig.1.) | Receiver's and Signaller's fitness respectively |
| --- | --- |
| 1, | $E_r = W(L, D) + r(V(L, D) - C(L, N))$ |
| | $E_s = V(L, D) - C(L, N) + rW(L, D)$ |
| 2, | $E_r = W(L, U) + r(V(L, U) - C(L, N))$ |
| | $E_s = V(L, U) - C(L, N) + rW(L, U)$ |
| 3, | $E_r = W(H, D) + r(V(H, D) - C(H, N))$ |
| | $E_s = V(H, D) - C(H, N) + rW(H, D)$ |
| 4, | $E_r = W(H, U) + r(V(H, U) - C(H, N))$ |
| | $E_s = V(H, U) - C(H, N) + rW(H, U)$ |
| 5, | $E_r = W(L, D) + r(V(L, D) - C(L, S))$ |
| | $E_s = V(L, D) - C(L, S) + rW(L, D)$ |
| 6, | $E_r = W(L, U) + r(V(L, U) - C(L, S))$ |
| | $E_s = V(L, U) - C(L, S) + rW(L, U)$ |
| 7, | $E_r = W(H, D) + r(V(H, D) - C(H, S))$ |
| | $E_s = V(H, D) - C(H, S) + rW(H, D)$ |
| 8, | $E_r = W(H, U) + r(V(H, U) - C(H, S))$ |
| | $E_s = V(H, U) - C(H, S) + rW(H, U)$ |

Before proceeding to the new set of solutions it is useful to recapture the conditions of honest signalling. Honest signalling under conflict of interest can be characterized by three sets of conditions [7]: (i) the receiver’s, (ii) the signaller’s (iii) and the conflict of interest condition. The receiver’s condition states that the receiver should react to different signaller differently. At the traditional signalling equilibrium it should give an Up (U) response to High quality signaller but it should turn Down (D) Low quality ones. Accordingly, the following inequalities must be fulfilled:

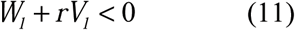

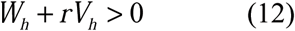

The signaller’s condition specifies that signallers should act differently at the honest equilibrium: High quality signallers should signal (S); low quality signallers should not signal (N) at the traditional signalling equilibrium. Accordingly, the potential benefits from signalling should be larger than the cost for high quality signallers and vice versa:

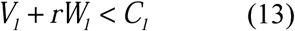

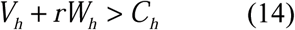

Last but not least, the conflict of interest should be specified. It implies that receiving the resource is beneficial for both signaller types:

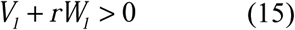

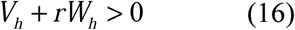

Note that in all of these conditions both the benefit and the cost denote differences between two actions (see Eqs. 5-10): giving or not giving the resource (*W\**), receiving or not receiving the resource (*V\**), and finally giving or not giving a signal (*C\**). Accordingly, *negative* values of *C*_*h*_ or *C*_*l*_ implies only that not giving a signal is more costly than giving (i.e. *C*(*\*,N*) > *C*(*\*,S*)); however, this condition tells nothing about the absolute values of *C*(*\*,S*) and *C*(*\*,N*). Here I investigate the possibility of negative cost (benefit) in the *absolute* sense, i.e. that both 0> *C*(*\*,S*) and 0> *C*(*\*,N*). Is honest signalling possible when signals for both types have benefits instead of costs?

## 3. Results

### Differential cost model

Since the conditions of honest signalling did not change, one have to check whether Eqs. 13 and 14 can be fulfilled alongside of the benefit assumption (i.e. 0> *C*(*\*,S*), *C*(*\*,N*)). substituting the cost functions (*C*(*\*,S*), *C*(*\*,N*)) into the equations we get:

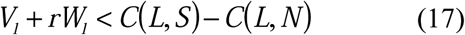

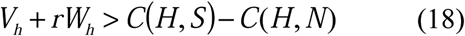

We can see, that in order for the first inequality to be satisfied the benefit from non-signalling has to be higher than the benefit from signalling for Low quality individuals; and it has to be higher so that non-signalling compensates Low quality signallers for the loss of not receiving the resource:

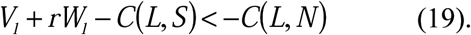

The opposite relation holds for High quality signallers:

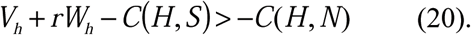

The benefit of non-signalling has to be smaller than the sum of the benefit they get receiving the resource and giving the signal.

Figures 2 and 3 depicts these and all other possible relations for Low and for High quality signallers respectively, in a differential cost model. There are five different regions for Low quality signallers (Fig. 2):

**Figure 2.**
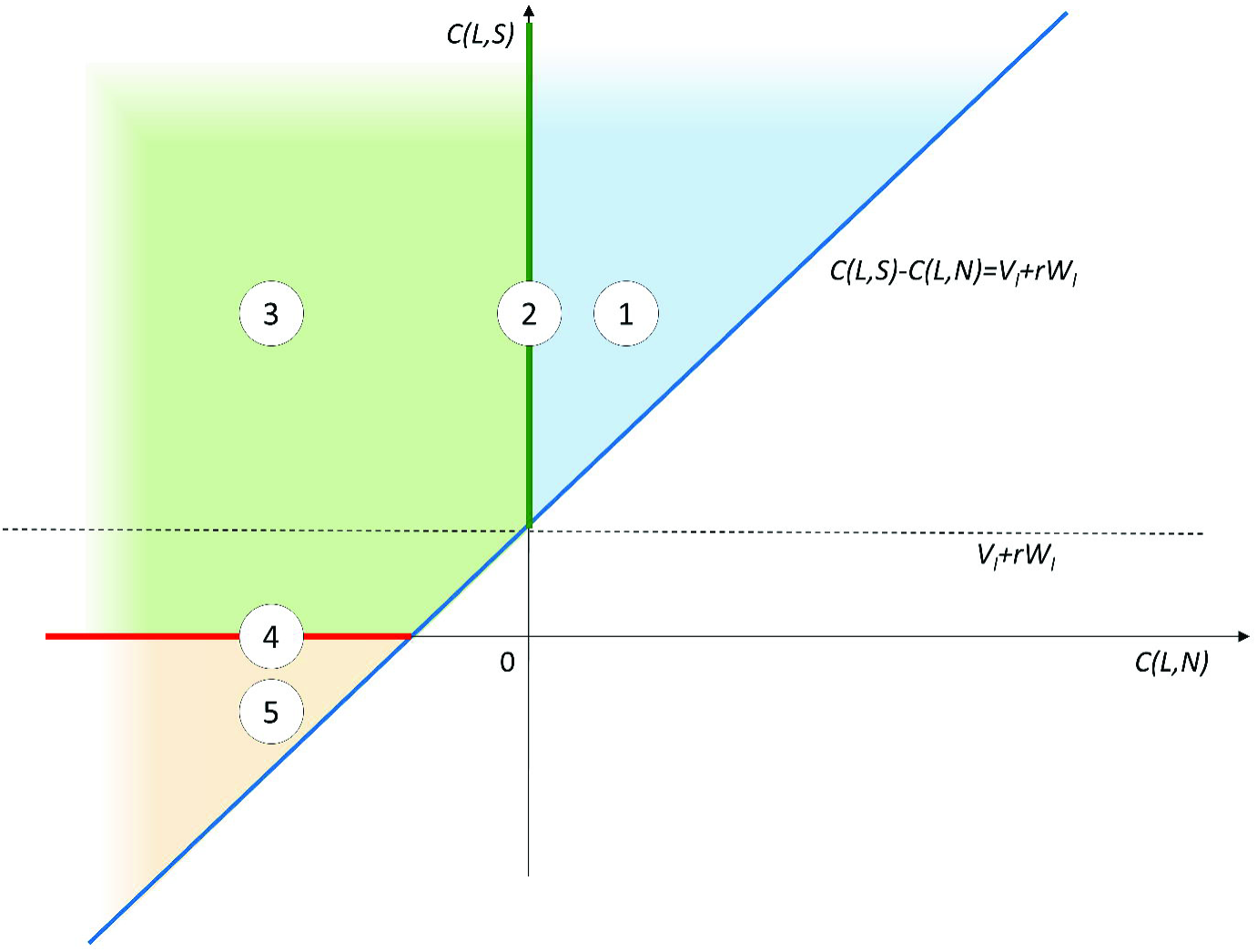
Regions where the difference between signalling and non-signalling for Low quality signallers allows honest signalling (i.e. it fits Eq 19) in differential cost models.

**Figure 3.**
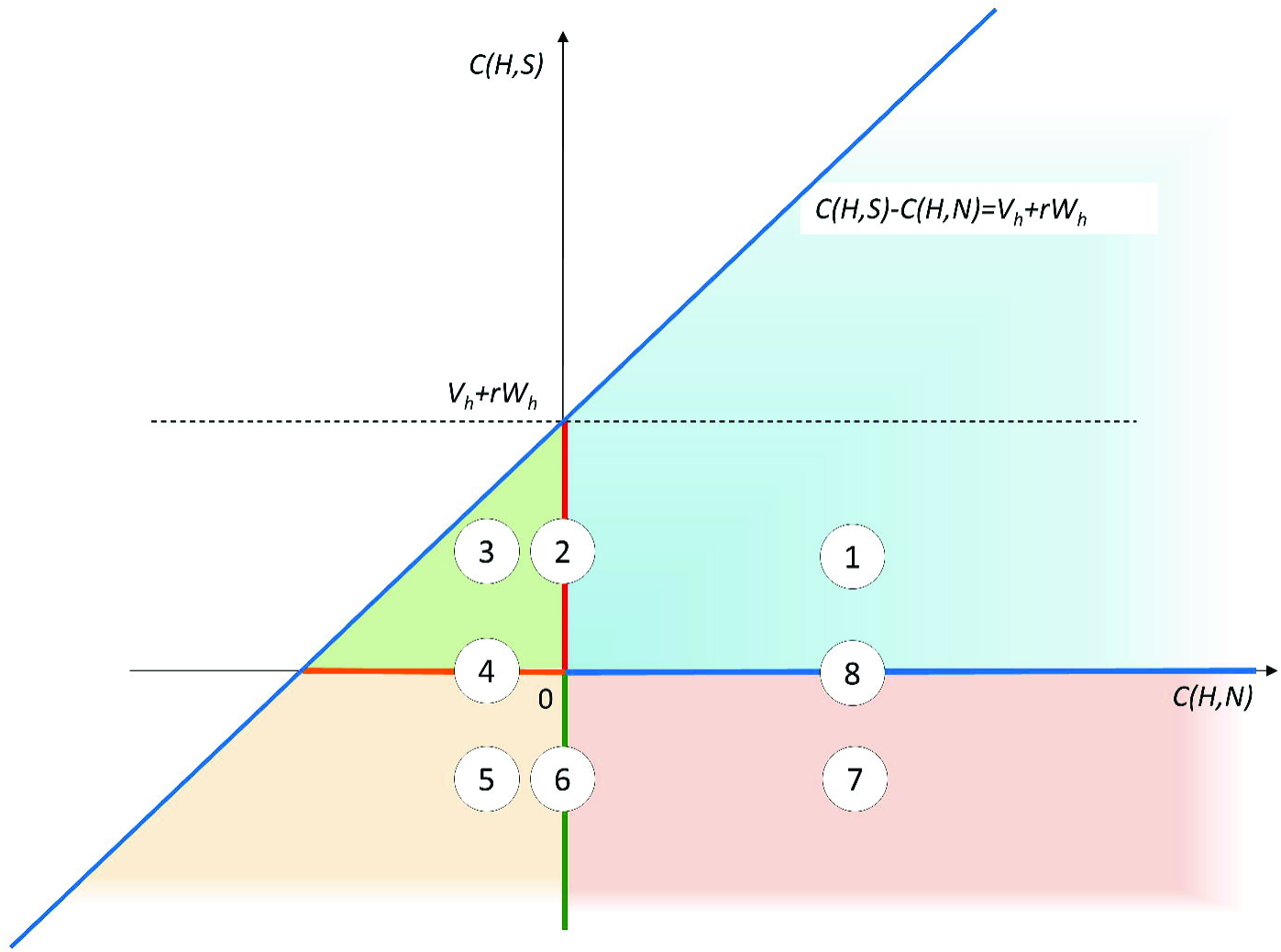
Regions where the difference between signalling and non-signalling for High quality signallers allows honest signalling (i.e. it fits Eq 20) in differential cost models.

i. in the first region both non-signalling (*C*(*L,N*)) and signalling (*C*(*L,S*)) is costly;
ii. in the second region (which denotes the line where *C*(*L,N*)=0) non-signalling has zero cost and signalling is costly, this is the standard set of assumptions of signalling models;
iii. in the third region non-signalling is beneficial (it has a negative cost) yet signalling is still costly;
iv. in the fourth region (which denotes the line where *C*(*L,S*)=0) non-signalling is beneficial and signalling has zero cost;
v. finally in the last, fifth region both non-signalling and signalling is beneficial. In other words, in this last region Low quality signallers get a benefit regardless of which action they chose, and this benefit is independent from the receiver’s response yet signalling still can be honest and evolutionarily stable.

Table 3 gives numerical examples for all regions (benefits in the model are as follows: *V*_*h*_ = 1, *V*_*l*_ = 1, *W*_*h*_= 1, *W*_*l*_ = −1).

**Table 3.** Numerical examples: differential cost model. Examples of *C*(*L,S*), *C*(*L,N*) are given for each region in Fig. 2. *C*_*l*_ = *C*(*L,S*)-*C*(*L,N*) = 1,2 in all regions (each example fits Eq. 19).

| Region | $C(L,S)$ | $C(L,N)$ |
| --- | --- | --- |
| 1, | 1.4 | 0.2 |
| 2, | 1.2 | 0 |
| 3, | 1 | -0.2 |
| 4, | 0 | -1.2 |
| 5, | -0.2 | -1.4 |

There are seven different regions for High quality signallers (Fig. 3):

i. in the first region both non-signalling (*C*(*H,N*)) and signalling (*C*(*H,S*)) is costly;
ii. in the second region (which denotes the line where *C*(*H,N*)=0) non-signalling has zero cost and signalling is costly;
iii. in the third region non-signalling is beneficial (it has a negative cost) yet signalling is still costly;
iv. in the fourth region (which denotes the line where *C*(*H,S*)=0) non-signalling is beneficial and signalling has zero cost;
v. in the fifth region both non-signalling and signalling is beneficial; (vi) in the sixth region signalling is beneficial yet non-signalling has zero cost;
vi. in the sixth region (which denotes the line where *C*(*H,N*)=0) non-signalling has zero cost and signalling is beneficial;
vii. in the seventh region non-signalling is costly, yet signalling is beneficial;
viii. and finally in the eights region (which denotes the line where *C*(*H,S*)=0) non-signalling is costly and signalling has zero cost.

Table 4 gives numerical examples for all regions (benefits are the same as before).

**Table 4.**
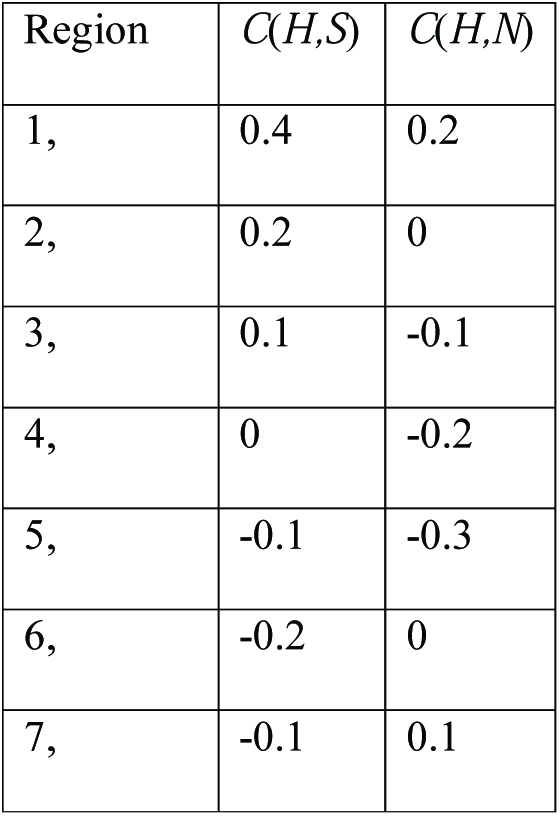
Numerical examples: differential cost model. Examples of *C*(*H,S*), *C*(*H,N*) are given for each region in Fig. 3. *C*_*h*_ = *C*(*H,S*)-*C*(*H,N*) = 0,2 except in region 6 and 7, where *C*_*h*_ = −0,2 (each example fits Eq. 20).

The traditional assumption is region 2 for both Low and High quality signallers (i.e. non-signalling has zero cost but signalling is costly). However, all these regions fit the conditions outlined in Eqs. 19 and 20 thus any combination of these regions is a solution. The important idea is that it is not a simple linear rescaling of the pay-offs for low and High quality signallers because these regions can be combined independently, which may result in unexpected or seemingly paradoxical parameter combinations that still can maintain honest signalling even under conflict of interest. All in all, there are 5x8=40 potential combinations; here I only discuss a few counter-intuitive examples.

(1) For example, it is possible that both non-signalling and signalling is costly for High quality signallers (Fig.3 region 1); yet both non-signalling and signalling is beneficial for Low quality signallers (Fig.2 region 5). In this example High quality signallers invest in signals and they are compensated by the receiver’s response, whereas Low quality signallers are compensated for the loss of receiver’s response by the benefit they receive for non-signalling.
(2) Interestingly enough the opposite is equally possible: that High quality signallers receive benefits for both non-signalling and signalling (Fig.3 region 5) yet Low quality signallers have to pay a cost for both non-signalling and for signalling (Fig.2 region 1). In this example signalling is costly for Low quality signallers which prevents them to mimic High quality ones, and High quality signallers receive an extra benefit on top of the receiver’s response.
(3) Perhaps the most interesting case where both Low and High quality signallers receive a benefit both from non-signalling and from signalling (region 5 in both Figs. 2 and 3). In this case there is no cost to signals in the system, everyone benefits from every single action, yet honesty still remains evolutionarily stable. In this example Low quality signallers are compensated for the loss of receiver’s response by the benefit they receive for non-signalling, whereas High quality signallers receive an extra benefit on top of the receiver’s response.

### Differential benefit model

What if the signal cost is the same for both types of signallers (i.e. we have a differential benefit model)? Is it still possible to get honest evolutionarily stable signalling with beneficial signals? The signaller’s conditions are modified as follows:

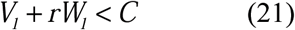

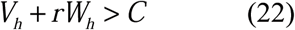

We can see that the same cost function has to satisfy both conditions. Accordingly, we have the following inequalities:

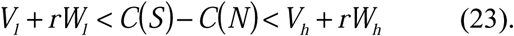

This implies that the difference between the costs of signalling and non-signalling has to be larger than the benefits from the Up response for Low quality signallers but this difference has to be smaller than the benefits from Up response for High quality signallers.

Figure 4 depicts the regions that satisfy the above condition in differential benefit models. There are five different regions in Fig. 4:

i. (i) in the first region both non-signalling (*C*(*N*)) and signalling (*C*(*S*)) is costly;
ii. (ii) in the second region non-signalling has zero cost and signalling is costly;
iii. (iii) in the third region non-signalling is beneficial (it has a negative cost) yet signalling is still costly;
iv. (iv) in the fourth region non-signalling is beneficial and signalling has zero cost;
v. (v) finally, in the last region both non-signalling and signalling is beneficial.

**Figure 4.**
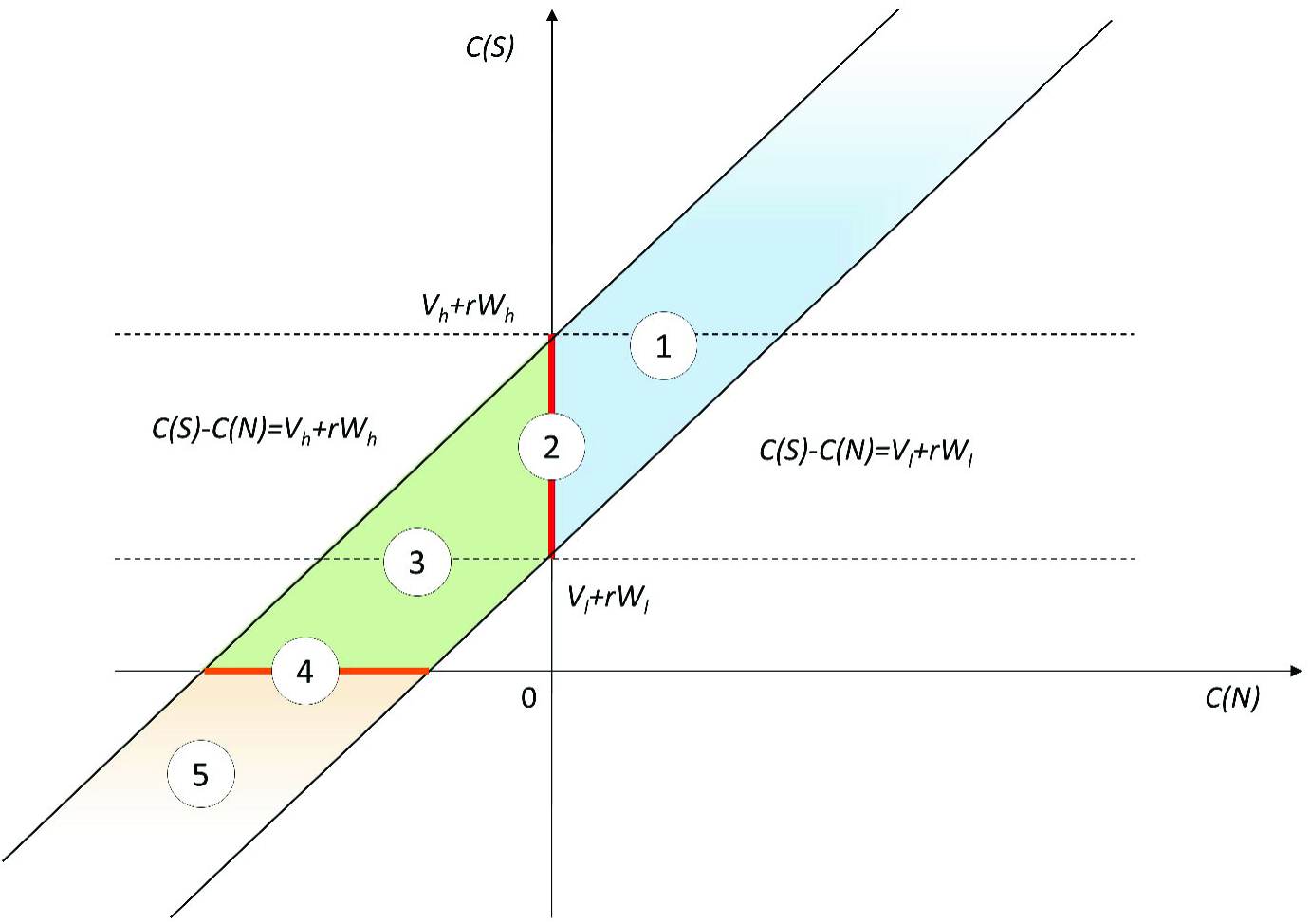
Regions where the difference between signalling and non-signalling allows honest signalling (i.e. it fits Eq 23) in differential benefit models.

The second region describes the traditional assumption of the signalling models and thus it corresponds to the classic Sir Philip Sydney game [15]. However, the most interesting is the fifth region, where just as before, signallers receive a benefit both from non-signalling and from signalling. Table 5 gives numerical examples for all regions (benefits in the model are as follows: *V*_*h*_ = 1, *V*_*l*_ = 0.5, *W*_*h*_ = 1, *W*_*l*_ = −1). Since signal cost is the same for Low and High quality individuals in differential benefit model thus changing the absolute value of cost corresponds to a linear rescaling in this case. However, the results show that this linear rescaling is possible (in any direction); it follows that the costly signalling equilibria of the ‘costly signalling’ models is a consequence of the *costly signalling assumption* (i.e. the choice of the second region, Fig. 2) and it is not a theoretical necessity.

**Table 5.**
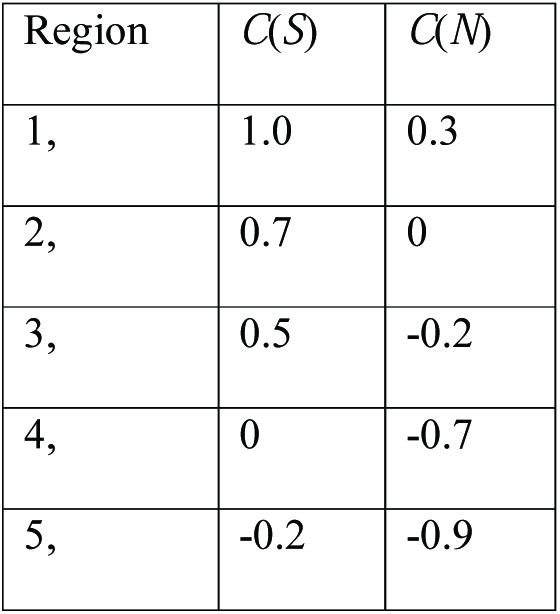
Numerical examples: differential benefit model. Examples of *C*(*S*), *C*(*N*) are given for each region in Fig. 4. *C* = *C*(*S*)-*C*(*N*) = 0,7 in all regions (each example fits Eq. 23).

## 4. Discussion

Here I showed that honest signalling needs no cost function. “Beneficial signals”, signals that have a benefit function instead of a cost function can maintain the honesty of communication. The conceptual importance of the model that it allows to separate signal cost (of any source) from the “potential cost of cheating”. It shows that signal cost - at or out of equilibrium - is not a condition of honest signalling. What maintains the honesty of communication is the potential cost of cheating, which is conceptually different from signal cost, as it can be a result of a benefit function. This “potential cost of cheating” is a fitness difference between two actions (to signal vs. not to signal) and this fitness difference can be negative even if both of the actions are beneficial on the first place.

Previous models were able to show that honest signals do not have to be costly for honest signallers to be evolutionarily stable, not even under conflict of interest [6-8]. The current result goes one step further, as it shows that signals need no cost at all to be honest. There is no need for production cost, maintenance cost, social cost, inclusive fitness cost, etc. This result invalidates Zahavi’s claim [4] about the special role of “signal selection”. Honest signalling is possible without signal cost: costly signalling is just one possible implementation, it is not a necessity.

The result also shows the limits of the ‘costly signalling’ paradigm [16, 17]. Costly signalling models in biology arrived at the conclusion of costly equilibrium because of the *costly signalling assumptions* of these models. In other words, the conclusion of the costly signalling models is built into the assumptions. Had the authors of these models investigated a benefit function instead of cost function, they would have arrived at the conclusion of beneficial equilibria. The ‘costly signalling’ assumption might be realistic and important, yet it is not a necessity or a ‘principle’.

Honest signalling and costly signalling have the same relation as natural selection vs. mendelian inheritance. Natural selection is the general principle: it assumes competition, reproduction, inheritance and variation. Mendelian inheritance is one possible implementation of an inheritance system that allows natural selection to work. Honest signalling is the general principle, costly signalling is a specific implementation that allows honest signalling to operate. Mendelian inheritance is not an overreaching “principle”, though it happens to be the most important inheritance system for “higher life”. The same way, “costly signalling” is not overreaching “principle” or necessity, though arguably it happens to be a very important mechanism of honest signalling.

Moreover, the Handicap Principle and the costly signalling paradigm is misleading because it suggested that measuring the “cost of signals” at the equilibrium provides valuable information about the source of honesty [1, 4]. As consequence hundreds of studies tried to measure out the equilibrium cost of signals without offering solid evidence in favour of the Handicap Principle [18, 19]. This is not surprising however, because measuring equilibrium cost is not informative, one has to measure out of equilibrium costs [8, 20]. However, measuring out of equilibrium cost in itself is not informative either. The cost is only informative in relation to the benefits of the action. What has to be measured is the pay-off resulting from the alternative actions (i.e. trade-offs). Unfortunately, the number of studies comparing out-of-equilibrium cost and benefits (i.e. signal trade-offs) is negligible (but see [21]).

All in all, signal cost is not a necessary ingredient of honesty: honesty needs no cost. Of course, it does not imply that signal cost cannot play a role in the maintenance of honesty; however, this is an empirical question and not a theoretical necessity.

## Author’s contributions

S.S. conceived the idea, analysed the model and wrote the article.

## Acknowledgements

This work was supported by National Research, Development and Innovation Office - NKFIH (OTKA) grant K 108974 and by the European Research Council (ERC) under the European Union’s Horizon 2020 research and innovation programme (grant agreement No 648693).

